# MULTIVALENT ACE2 ENGAGEMENT ACTIVATES PIEZO1–SPHINGOMYELINASE SIGNALING IN SARS-COV-2 SPIKE-MEDIATED CELL–CELL FUSION

**DOI:** 10.64898/2026.10.22.761047

**Authors:** Elena M. Zaitseva, Eugene N. Zaitsev, Kamran Melikov, Leonid V. Chernomordik

## Abstract

SARS-CoV-2 Spike not only fuses the viral envelope with host membranes to deliver viral RNA but also drives syncytium formation, a process that may contribute to viral spread and pulmonary pathology. Although the conformational transitions in Spike triggered by ACE2 engagement are well characterized, the host-cell signaling events initiated by Spike–ACE2 interactions remain less understood. Here, we examine how Spike activates acid sphingomyelinase (ASM) in ACE2-expressing A549 cells and how this activity promotes cell–cell fusion. Using ASM-deficient and complemented cells, exogenous ceramide, pharmacological perturbations, and a split-NeonGreen fusion assay, we support a fusion-promoting pathway in which Spike-induced ACE2 clustering activates the mechanosensitive ion channel PIEZO1, triggering lysosomal exocytosis. Secreted ASM, together with intracellular neutral sphingomyelinase, generates ceramide that enhances syncytium formation, likely through activation of protein phosphatases PP1 and PP2A. Our results suggest that these phosphatases promote fusion by facilitating actin cortex detachment and reducing actomyosin tension through dephosphorylation of ERM proteins. Our findings highlight dynamic changes in membrane lipid composition and membrane–cytoskeleton interactions as regulators of fusion and signaling in SARS-CoV-2 infection and, potentially, other viral and cellular systems.

## INTRODUCTION

Cell–cell fusion is a biological process in which two or more cells merge their plasma membranes and combine their cytoplasmic contents to form a single, multinucleated cell. This process is essential for development, immunity, tissue repair, and certain diseases, and many enveloped viruses exploit it using fusogenic proteins to induce fusion between infected and neighboring cells [1]. SARS-CoV-2 similarly promotes syncytium formation in the lungs through spike protein– mediated membrane fusion. Syncytia were detected in lung tissue from 18 of 24 deceased patients examined and suggested to facilitate immune evasion and direct cell-to-cell viral spread [2]. Syncytium formation and growth may contribute to pulmonary pathology, including impaired vascular barrier function, although this possibility remains to be established.

SARS-CoV-2 Spike-mediated membrane fusion is driven by conformational changes in Spike triggered by its interaction with the ACE2 receptor [3]. However, fusion also depends on the lipid composition of the interacting membranes. Phosphatidylserine, cholesterol, phosphatidylethanolamine, and ceramide, lipids that modulate membrane fluidity, curvature, membrane-domain organization, and the spatial distribution of viral and cellular fusion factors [3, 4], have all been implicated as positive regulators of fusion [3–5]. Ceramide, one of the lipids discussed as a potential modulator of different fusion processes (reviewed in [6]) is generated by ASM and neutral sphingomyelinase (nSMase2). While steady-state ceramide levels in the plasma membrane are regulated primarily by nSMase2 [7], ASM is considered a major source of ceramide generated in the outer leaflet of the plasma membrane in response to cellular stress [8]. ASM is a lysosomal enzyme that can be released to the cell surface by Ca²⁺-dependent lysosomal exocytosis and plays important roles in plasma membrane repair and remodeling, immune responses, and inflammation [9, 10].

Several studies document SARS-CoV-2-induced translocation of ASM from lysosomes to the plasma membrane and suggest an important role for the sphingomyelinase–ceramide pathway in viral infection [11–13]. It has been suggested that ASM-generated ceramide in the outer leaflet of the plasma membrane spontaneously clusters into small and then larger ceramide-enriched gel-like platforms that facilitate viral entry by promoting the clustering of ACE2, the cellular receptor for SARS-CoV-2 [10, 11, 13, 14]. However, since spike–ACE2 engagement has already occurred, the clustering of ACE2 may not be the primary purpose of ceramide platforms, although it may contribute to subsequent stages of signaling and membrane remodeling. Consistent with this, viral fusion and entry were found in [15] to occur independently of ACE2 localization in lipid rafts, suggesting alternative mechanisms that may involve receptor clustering, mechanical perturbation, co-receptor engagement, or stress-response pathways that induce lysosomal exocytosis [16, 17]. While the dependence of SARS-CoV-2 entry and infection on ASM and ceramide is well documented, the role of ASM in Spike-mediated fusion of ACE2-expressing cells remains unexplored. Moreover, the mechanisms by which Spike induces ASM translocation to the surface of ACE2-expressing cells, and whether and how ASM-generated ceramide promotes cell–cell fusion, remain unknown.

In the present manuscript, we propose a comprehensive model of the signal transduction pathway linking Spike–ACE2 receptor engagement to cytoskeletal remodeling in target cells. This model integrates membrane-associated signaling, lipid metabolism, and cytoskeletal dynamics to explain how Spike–ACE2 interactions may translate extracellular viral recognition into intracellular processes that promote membrane remodeling and potentially facilitate Spike-mediated cell–cell fusion. Our findings support a model in which ceramide production serves chiefly as an outside-in signaling conduit linking ACE2–Spike interaction to cytoskeletal reorganization. This proposed mechanism contrasts with the widely accepted model, in which ceramide platforms primarily increase receptor concentration and function as structural facilitators of viral fusion.

## RESULTS

### ACE2 ligation and PIEZO1 activation increase ASM externalization

To explore the effects of Spike-ACE2 interactions on ACE2-expressing cells, we used A549 cells, a human alveolar basal epithelial adenocarcinoma cell line widely used as an *in vitro* model for SARS-CoV-2/COVID-19, lung cancer, and alveolar type II cell research [18, 19]. We transduced A549 cells to express ACE2 under the control of a weak SV40 promoter to avoid ACE2 levels substantially exceeding relatively low physiologically relevant receptor densities, such as those found in human lung epithelium tissue [20]. In addition, limiting ACE2 expression in our A549-ACE2 cells reduced the likelihood of spontaneous ACE2-dependent signaling events, which we observed in our preliminary experiments when ACE2 was strongly overexpressed. In another consideration, A549 cells were selected as the parental cell line because of their reported low cell-surface expression of TLR2 and TLR4 under basal conditions [21]. SARS-CoV-2 Spike has been reported to activate cell-surface TLRs, particularly TLR2 [22] and, more controversially, TLR4 [23, 24] stimulating downstream inflammatory pathways independent of Spike-ACE2 interactions. Thus, using A549 cells allowed us to minimize potential contributions of TLR signaling.

Because the ACE2 cytoplasmic tail lacks canonical signaling capacity, Spike–ACE2 engagement is thought to trigger intracellular responses indirectly rather than through a traditional receptor-signaling mechanism [25]. Spike–ACE2 interaction has been reported to induce ACE2 clustering [14, 26]. We reasoned that clustering of membrane proteins can generate membrane deformation [27] and induce signaling by opening a mechanically activated Ca^2+^ channel, PIEZO1 [28], already implicated in COVID-19 pathogenesis [29–31]. In this framework, Spike-induced ACE2 clustering may trigger mechanotransduction, linking receptor engagement to PIEZO1 activation and downstream Ca²⁺-dependent signaling.

To examine the role of PIEZO1 in lysosomal exocytosis leading to ASM release, we established a cell-based reporter assay in A549-ACE2-ASM-KO cells stably transduced with a lentiviral construct encoding ASM-pHluorin protein. This pH-sensitive fluorescent reporter enables detection of extracellular ASM by monitoring an increase in pHluorin fluorescence after release into the neutral extracellular environment. To enhance detection of this relatively weak signal, we incorporated a GFP-specific fluorescently labelled nanobody (GFP-Booster 647) into the assay.

Application of PIEZO1 agonist Yoda2 [32] to A549-ACE2-ASM-KO cells stably expressing ASM-pHluorin resulted in its externalization, detected as an increased pHluorin fluorescence (Fig. 1A,B). Importantly, replacing Yoda2 with soluble trimeric SARS-CoV-2 Spike protein (10 µg/mL) induced a similar increase in fluorescence, indicating similar level of ASM-pHluorin externalization. The hypothesis that Spike-induced ASM-pHluorin externalization involves PIEZO1 was supported by its inhibition by GsMTx4, an inhibitor of mechanosensitive channels, including PIEZO1 [33].

**Figure 1.**
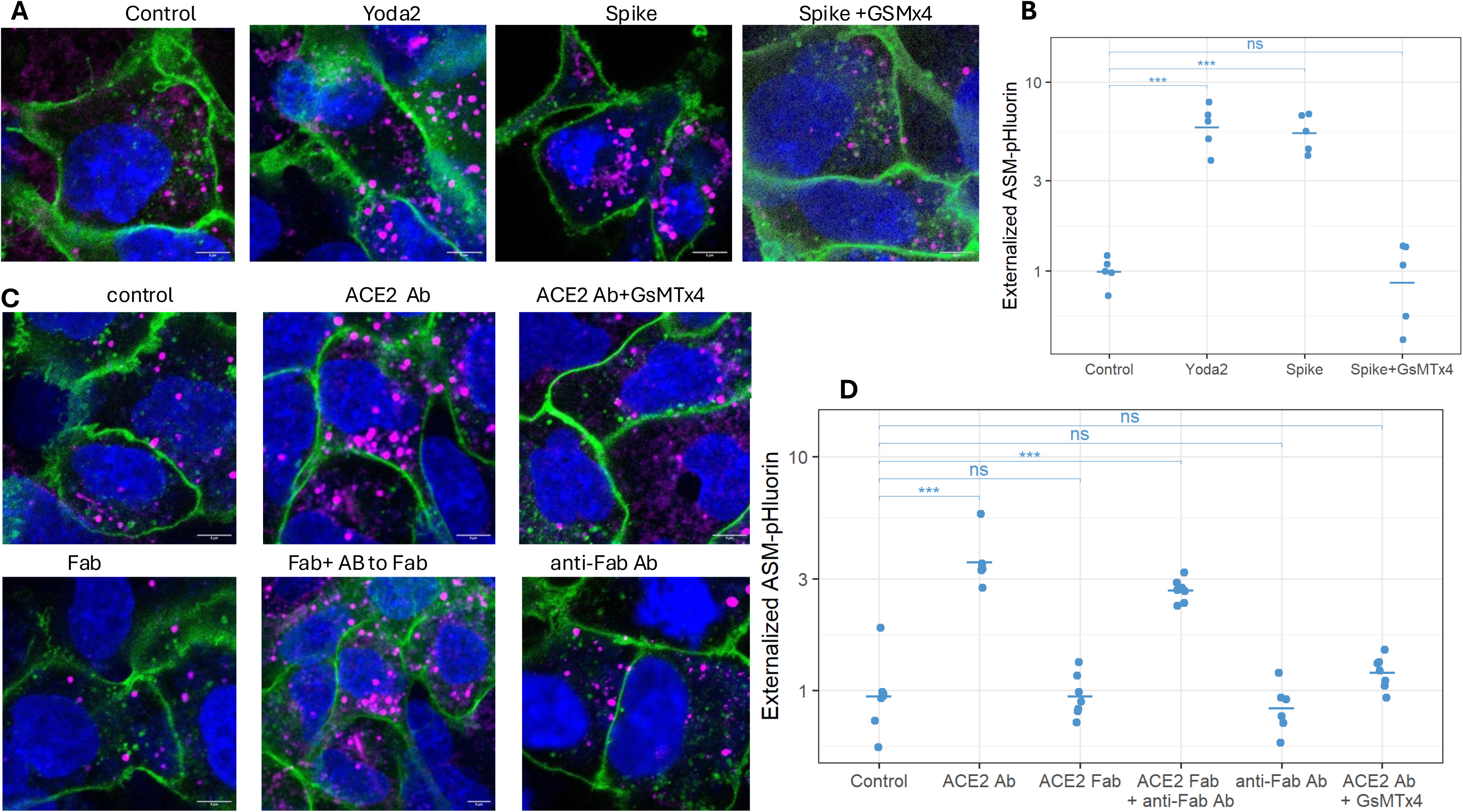
Spike and multivalent ACE2 ligation increase ASM-pHluorin surface signal. (A, B) A549-ACE2 *-*ASM-KO cells expressing ASM-pHluorin were incubated for 4 hours at room temperature with soluble trimeric Spike (10 µg/mL), Yoda2 (5 µM), or Spike plus GsMTx4 (12 µM). GFP-Booster 647 staining was quantified as described in the Methods. (C, D) Cells were treated with ACE2 antibody (20 µg/mL), Fab (7 µg/mL), or Fab plus anti-Fab antibody (30 µg/mL) or anti-Fab antibody alone; indicate whether GsMTx4 was included in the plotted experiment. (A,C) Representative fields for the respective conditions. Scale bar, 5 µm. In (B,D), points represent independent experiments, with the horizontal bar indicating the group mean. Statistical significance of pairwise comparisons is denoted by asterisks (*p < 0.05, **p < 0.01, ***p < 0.001; ns, not significant)

To model ACE2 ligation by the Spike protein, cells were treated with either an ACE2-specific antibody or the corresponding Fab fragment. Bivalent ACE2 ligation by the antibody—but not monovalent ligation by the Fab fragment—strongly induced ASM externalization. Co-treatment with the Fab fragment and an anti-Fab antibody restored bivalent ligation and ASM externalization. This response also depended on mechanosensitive channels, as GsMTx4 abolished ASM externalization (Fig.1C,D).

Because ASM deficiency may affect multiple physiological processes and potentially introduce experimental artifacts, we modified our reporter assay to directly assess lysosomal exocytosis independently of ASM. Specifically, we used LAMP1-pHluorin, a well-established lysosomal membrane marker, to monitor lysosomal exocytosis in A549 WT and A549 ASM-KO cells. Notably, ASM deficiency did not impair lysosomal exocytosis in A549 ASM-KO cells, as assessed by LAMP1-pHluorin fluorescence (Fig.2A,B). These findings indicate that loss of ASM does not intrinsically disrupt the lysosomal exocytosis machinery in this cellular model.

**Figure 2.**
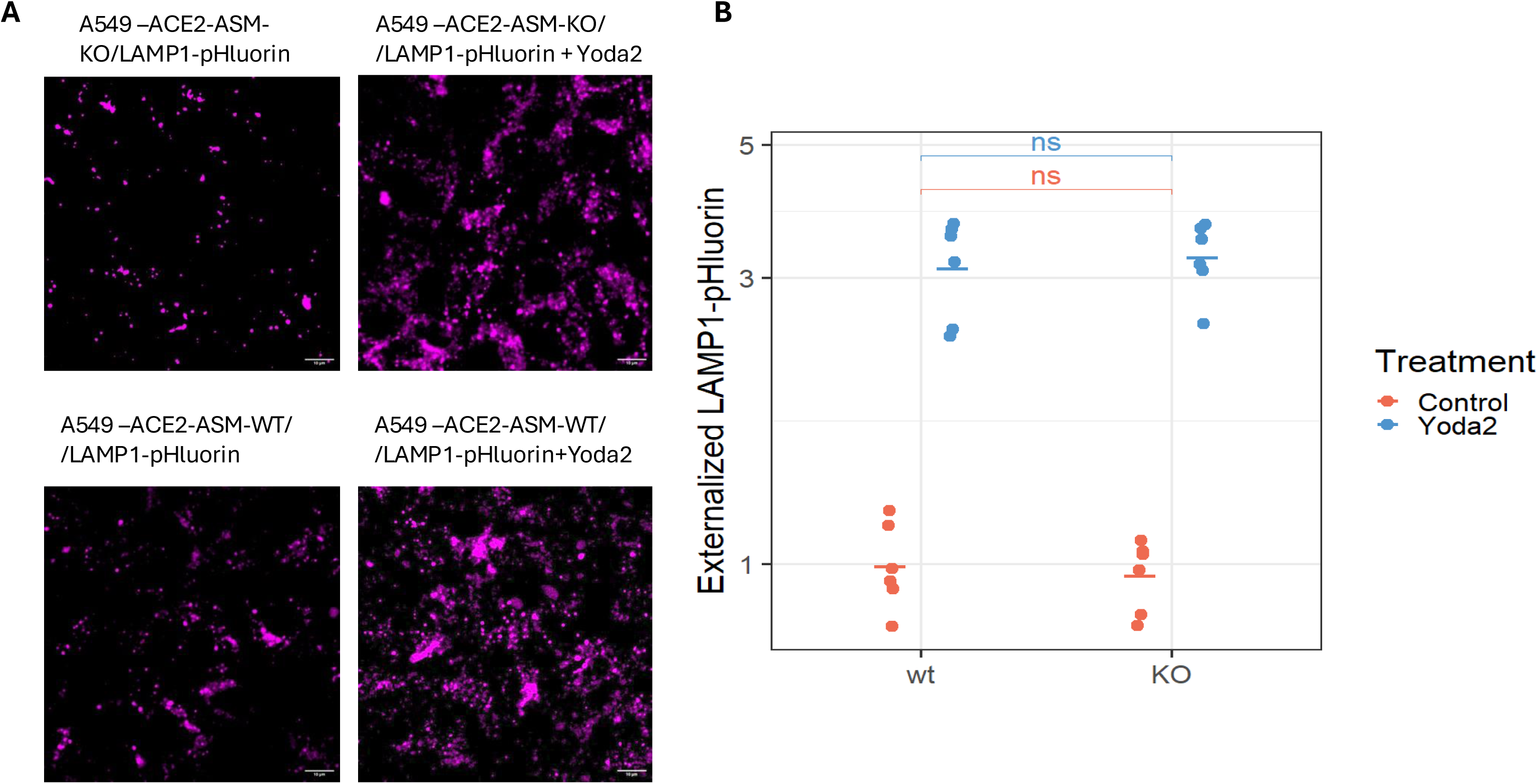
ASM loss does not eliminate the Yoda2-induced LAMP1-pHluorin response. (A). Representative images of ACE2-expressing A549-ACE2-ASM-WT (wild-type) and A549-ACE2-ASM-KO cells (both lines stably expressing LAMP1-pHluorin) were untreated or treated with Yoda2 (5 µM) for 4 hours at room temperature. Accessible reporters were stained with GFP-Booster 647. Scale bar, 10 µm. (B). The images were analyzed as in Fig.1. Points represent independent experiments, with the horizontal bar indicating the group mean. Statistical significance of pairwise comparisons is denoted by asterisks (*p < 0.05, **p < 0.01, ***p < 0.001; ns, not significant)

Together, our findings indicate that ACE2-Spike protein interactions promote ASM externalization via lysosomal exocytosis in ACE2-expressing cells through a pathway that depends on ACE2 ligation and PIEZO1 activation.

### ASM and ceramide are cofactors in Spike-mediated cell–cell fusion

To assess the role of ASM and ceramide in Spike protein–mediated cell–cell fusion, we developed a cell–cell fusion assay based on a split monomeric NeonGreen fluorescent protein system, as previously described [34]. We co-plated HEK293 cells stably expressing the SARS-CoV-2 Spike protein along with x5NG11 fragment and A549-ACE2 cells expressing mNG3K1-10. Cell fusion brought together the two complementary fragments of the split NeonGreen protein to reconstitute a functional fluorescent protein. The resulting fluorescence signal therefore provided a quantitative readout of Spike protein–mediated cell–cell fusion. We compared the fusion efficiency of A549-ACE2-ASM-WT (wild-type) and A549-ACE2-ASM-KO acceptor cells and found that ASM knockout resulted in an approximately 3-fold reduction in fusion activity compared with ASM WT cells (Fig.3A,B). Both exogenous ceramide supplementation and genetic ASM complementation largely restored fusion activity in ASM-deficient cells (A549-ACE2-ASM-KO), whereas their effects on fusion in ASM-WT (A549-ACE2-ASM-WT) cells were significant but much less pronounced. Activation of the PIEZO1 channel with YODA2 agonist significantly enhanced fusion activity in both A549-ACE2-ASM-WT cells and in A549-ACE2-ASM-KO cells genetically complemented with ASM (A549-ACE2-ASM-KO+ASM cells); however, YODA2 failed to increase fusion in A549-ACE2-ASM-KO cells (Fig.3A,B). These findings suggest that PIEZO1 acts upstream of ASM in promoting cell–cell fusion, and that PIEZO1 activation enhances fusion primarily by driving ASM externalization.

**Figure 3.**
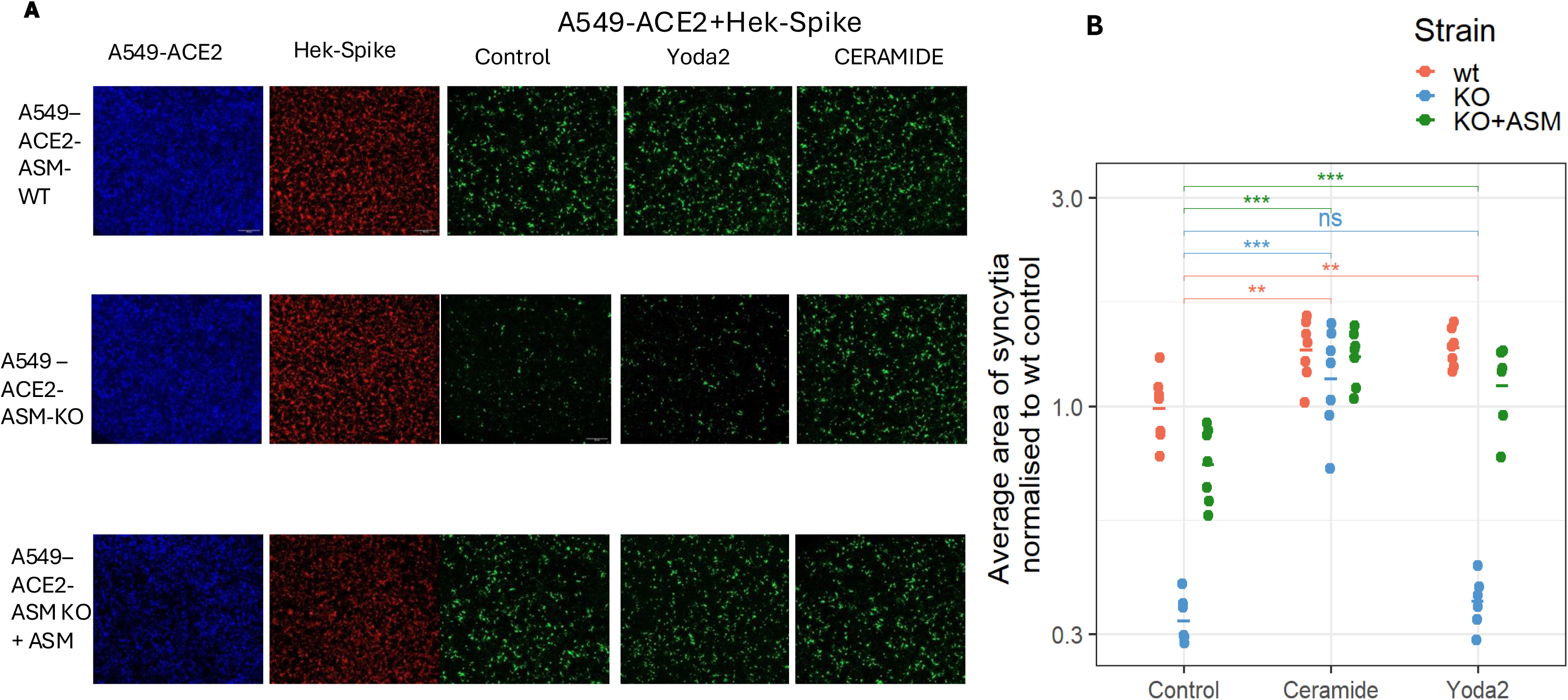
ASM and exogenous ceramide increase Spike-mediated cell–cell fusion. (A) Representative mNeonGreen images of Spike-expressing HEK293T/17 cells co-cultured with matched A549-ACE2-ASM-WT, A549-ACE2-ASM-KO or A549-ACE2-ASM-KO genetically complemented target KO cells, with or without Yoda2 (5 µM) or ceramide (10 µM). Co-cultures were incubated for 2 hours at room temperature to facilitate heterotypic cell contacts, followed by 36 hours at 37 °C. Scale bar, 500 µm. (B) Average areas of syncytia were normalized to wild-type control and plotted on a log₁₀ scale. Points represent independent experiments, with the horizontal bar indicating the group mean. Statistical significance of pairwise comparisons is denoted by asterisks (*p < 0.05, **p < 0.01, ***p < 0.001; ns, not significant).

### The Role of nSMase in Promoting Cell–Cell Fusion

The major sphingomyelinases in cells are acid sphingomyelinase (ASM) and neutral sphingomyelinase (nSMase). These enzymes generally operate at different leaflets of the plasma membrane and represent stimulus-responsive sources of distinct pools of ceramide that converge on membrane remodeling. Whereas ASM activation is frequently associated with lysosomal stress and lysosomal exocytosis [9], nSMase—particularly nSMase2—is activated by receptor-associated and plasma-membrane stress signals [35]. Thus, Spike–ACE2 engagement may recruit these sphingomyelinase pathways through distinct upstream mechanisms, with their relative contributions depending on cellular context, subcellular localization, and the nature and duration of the stimulus. Importantly, ASM and nSMase need not function as independent pathways; previous studies suggest that they can act sequentially or in parallel, with activation of one pathway influencing the activity of the other [36, 37].

Accordingly, we assessed whether nSMase contributes to Spike-mediated cell–cell fusion by pharmacologically inhibiting its activity during the fusion assay. In fusion of Spike-expressing HEK293 cells with A549-ACE2-ASM-KO cells, the nSMase inhibitors DPTIP [38] and nSMase2-IN [39] decreased fusion by only ∼15%. In contrast, when acceptor cells expressed ASM (A549-ACE2-ASM-WT cells), the same nSMase inhibitors produced a much stronger, ∼50% decrease in fusion (Fig. 4). These findings indicate that nSMase contributes to Spike-mediated cell–cell fusion and, interestingly, that this contribution is substantially greater in the presence of functional ASM. The differential effect of nSMase inhibition under ASM-deficient versus ASM-active conditions further suggests functional interplay between ASM- and nSMase-dependent pathways in regulating ceramide availability and cell–cell fusion.

**Figure 4.**
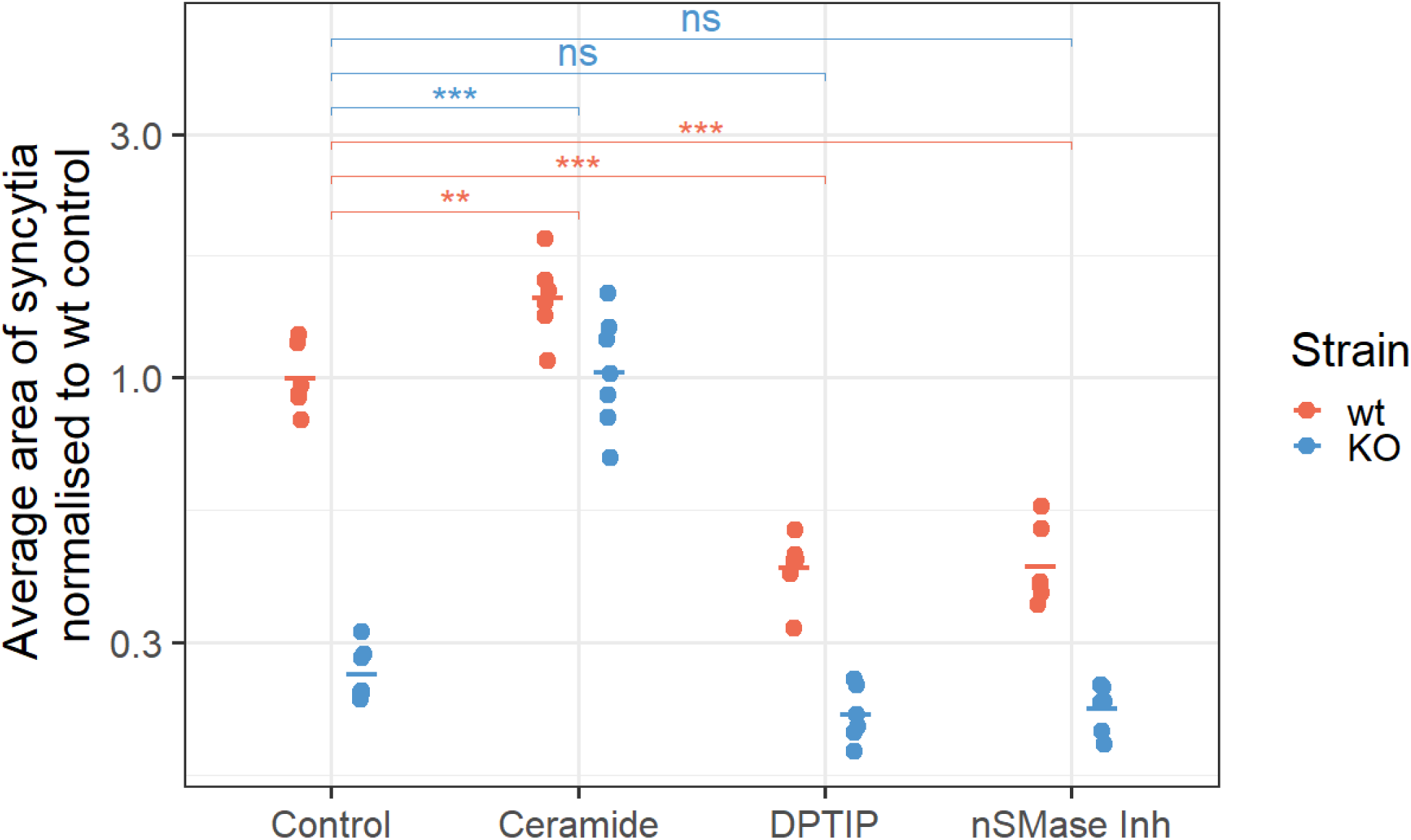
Effect of putative nSMase inhibitors on cell-cell fusion differs by ASM status of the target cells. Spike-expressing HEK293T/17 cells were co-cultured with A549-ACE2-ASM-WT or A549-ACE2-ASM-KO cells in the presence of ceramide (10 µM) or the inhibitors of nSMase: DPTIP and nSMase2-IN1 (10 µM each), where indicated. Average areas of syncytia were normalized to wild-type control and plotted on a log₁₀ scale. Points represent independent experiments, with the horizontal bar indicating the group mean. Statistical significance of pairwise comparisons is denoted by asterisks (*p < 0.05, **p < 0.01, ***p < 0.001; ns, not significant)

### Activation of PP1 and PP2A protein phosphatases and inactivation of ERM proteins

identified a pathway linking Spike–ACE2 interaction to ASM externalization and, presumably, to increased ceramide, we next considered possible mechanisms by which elevated ceramide may promote fusion. Ceramides are bioactive lipids that function as signaling molecules and influence diverse cellular processes, including apoptosis, cell-cycle arrest, and differentiation [40]. In addition to altering the biophysical properties of lipid bilayers [40], ceramides can act through interactions with ceramide-binding proteins, which are localized at the plasma membrane and in intracellular compartments [41]. Among the best-characterized ceramide-responsive proteins are the protein phosphatases PP1 and PP2A, which regulate the phosphorylation state of numerous cellular substrates. Ceramide has been reported to activate PP1 and PP2A [42], thereby promoting dephosphorylation of downstream target proteins.

We reasoned that if the pro-fusion activity of ceramide involves activation of PP1 and PP2A, then direct pharmacological activation of these phosphatases might recapitulate part of the downstream signaling induced by ceramide. Indeed, treatment of A549-ACE2-ASM-KO cells fusing with Spike-expressing HEK293 cells with pharmacological activators of PP1-SMAPP1 [43] and the PDP3 peptide [44] or of the proposed PP2A modulator, DT-061 [45] substantially rescued the fusion defect associated with ASM deficiency (Fig. 5).

**Figure 5.**
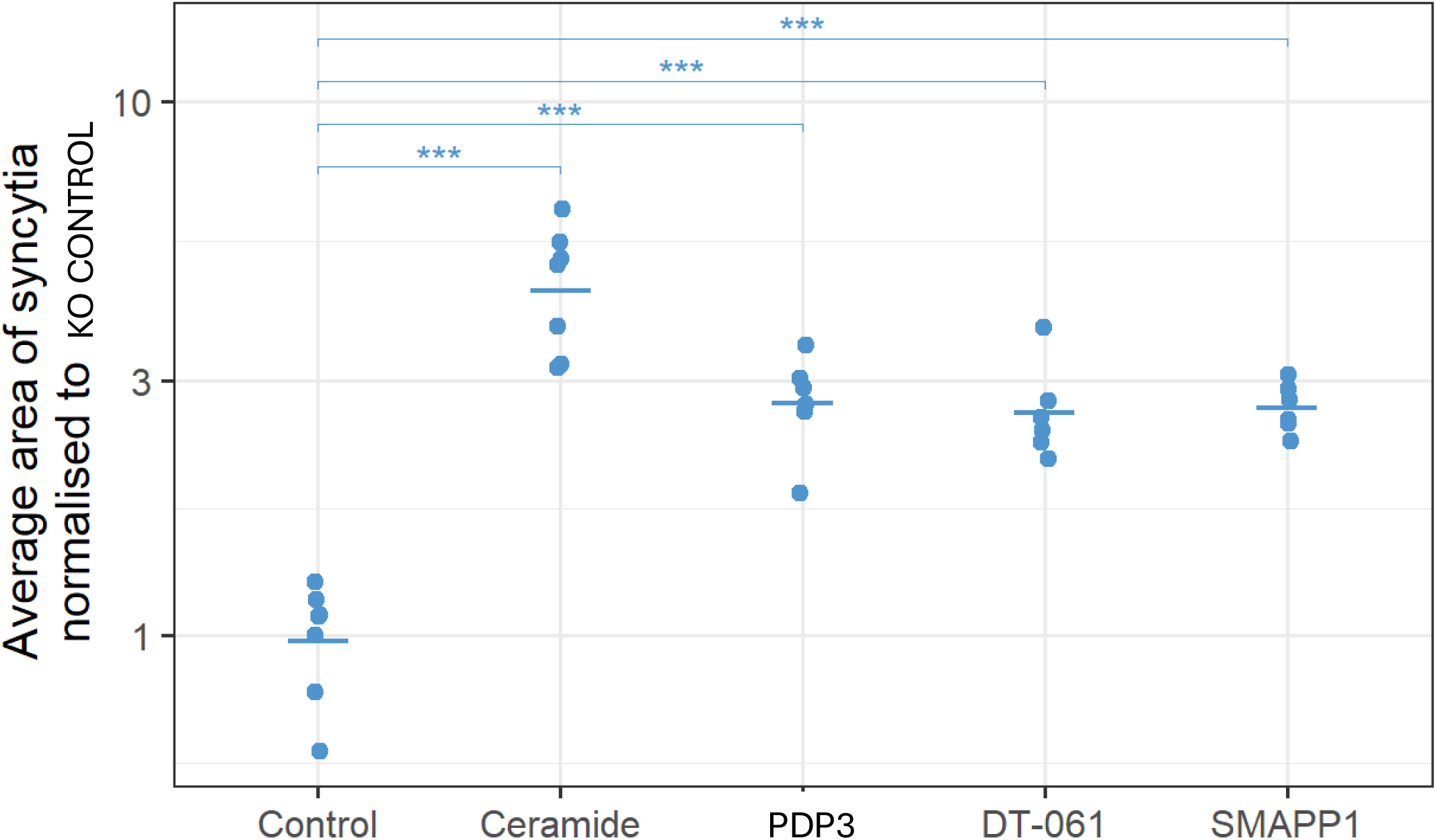
PP1/PP2a -related compounds increase cell-cell fusion for ASM-knockout target cells. A549-ACE2-ASM-KO cells were exposed to SMAPP1, DT-061, ceramide (10 µM for each), or PDP3 (12 µM) during co-culture with Spike-expressing HEK293T/17 cells. Average areas of syncytia were normalized to wild-type control and plotted on a log₁₀ scale. Points represent independent experiments, with the horizontal bar indicating the group mean. Statistical significance of pairwise comparisons is denoted by asterisks (*p < 0.05, **p < 0.01, ***p < 0.001; ns, not significant)

PP1 and PP2A dephosphorylate numerous cellular substrates, including proteins of the ERM (ezrin, radixin, and moesin) family [46, 47]. ERM proteins function as cytoskeletal linkers that connect the plasma membrane to the underlying actin cortex [48]. Their activation involves binding to phosphatidylinositol 4,5-bisphosphate (PIP2) at the plasma membrane, which promotes conformational activation and phosphorylation. Activated ERM proteins strengthen membrane–actin cortex attachment, thereby helping to maintain and organize the cortical cytoskeleton. Conversely, ERM dephosphorylation and/or depletion of PIP2 can promote ERM inactivation and weakening of membrane–cortex attachment. PIP2 depletion can occur through PLC-mediated hydrolysis or inhibition of PI4-kinase activity.

Previous studies have reported that ASM activation or exogenous ceramide treatment can promote ERM and actin-cortex detachment through PP2A-dependent mechanisms [47]. Detachment of the cortical actin network has also been implicated in Spike-mediated cell–cell fusion [49], and reported to promote the cell–cell fusion stage of multinucleated osteoclast formation [50–52].

We therefore asked whether the ACE2/PIEZO1/ASM/ceramide/phosphatase pathway could promote Spike-mediated cell fusion by weakening membrane–cortex attachment. As shown in Fig.6, two interventions that reduce PIP2 levels—the PLC activator m-3M3FBS [53] and the PI4KA inhibitor GSK-A1 [54] —as well as two ezrin inhibitors, NSC305787 and MMV667492 [55], partially rescued the fusion defect observed when Spike-expressing HEK293 cells were fused with A549-ACE2-ASM-KO cells. Blebbistatin, an inhibitor of non-muscle myosin II that reduces actomyosin contractility and cytoskeletal tension [56], also partially rescued the fusion defect (Fig. 6). This result is consistent with previous findings implicating actomyosin contractility in the regulation of cell–cell fusion [49] and suggests that excessive cortical tension may contribute to the fusion defect in ASM-KO cells.

**Figure 6.**
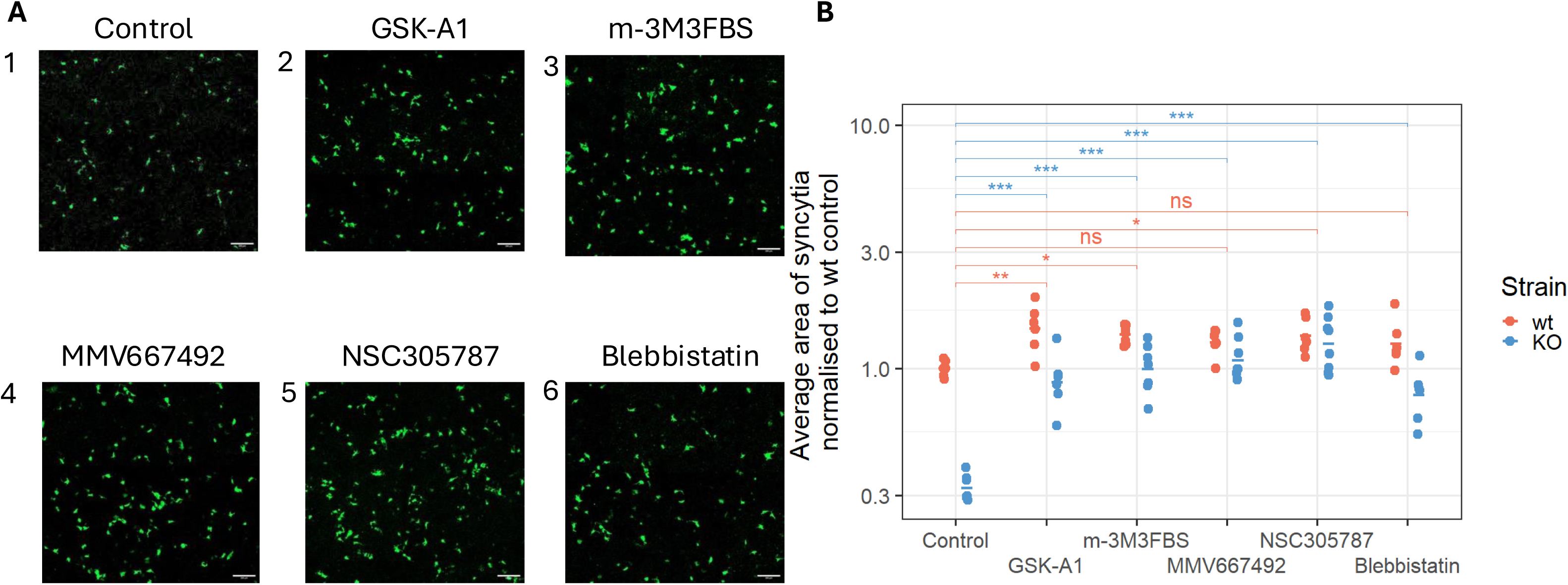
Compounds that affect cortical organization increase cell-cell fusion. Spike-expressing HEK293T/17 cells were co-cultured with A549-ACE2-ASM-WT or A549-ACE2-ASM-KO cells in the presence of GSK-A1 (10 µM), m-3M3FBS (10 µM), MMV667492 (5 µM), NSC305787 (10 µM), or blebbistatin (10 µM) for 2 hours at RT, followed by 36 hours at 37 °C. (A) Representative images from randomly chosen fields of co-culture knockout cells with or without treatment. Scale bar, 200 µm (B) Average areas of syncytia were normalized to wild-type control and plotted on a log₁₀ scale. Points represent independent experiments, with the horizontal bar indicating the group mean. Statistical significance of pairwise comparisons is denoted by asterisks (*p < 0.05, **p < 0.01, ***p < 0.001; ns, not significant).

Notably, some of the same pharmacological perturbations (GSK-A1, m-3M3FBS and NSC305787) modestly enhanced syncytium formation in ASM-WT cells. That finding suggests that membrane–cortex attachment and actomyosin contractility constitute general constraints on fusion and that the balance between kinase and phosphatase activities regulating ERM may control the fusogenicity of the cell membrane. It also suggests that the signaling capacity is not fully saturated and further modulation of PP1/PP2A activity and ERM function can enhance fusion even in cells with functional ASM.

Thus, ERM-dependent membrane–cortex attachment and actomyosin contractility appear to represent broader regulatory constraints on Spike-mediated fusion rather than processes operating exclusively downstream of ASM.

Collectively, these findings, and, especially, the finding that the reagents that weaken ERM-dependent membrane–cortex attachment or reduce actomyosin contractility have much stronger effects on fusion of ASM-deficient target cells (Fig. 6), are consistent with a model in which ASM/ceramide-dependent signaling contributes to establishing a membrane and cytoskeletal state that is permissive for efficient fusion. However, the pharmacological rescue experiments do not, by themselves, establish that ERM proteins constitute an obligatory linear downstream pathway of ASM/ceramide signaling. Rather, they support a model in which ASM/ceramide, phosphatase activity, ERM-dependent membrane–cortex attachment, and actomyosin contractility converge to regulate the cellular state required for efficient membrane fusion.

## DISCUSSION

In this paper, we explored how interactions of the SARS-CoV-2 Spike protein with the ACE2 receptor activate changes in lipid metabolism in ACE2-expressing cells to increase the efficiency of cell-cell fusion. Based on our findings, we propose a pathway in which Spike-induced ACE2 clustering and subsequent PIEZO1 channel activation promote lysosomal exocytosis. Secreted ASM, acting in concert with nSMase, generates ceramide that promotes Spike-mediated cell-cell fusion by mechanisms that may involve activation of the protein phosphatases PP1 and PP2A, which promote actin cortex detachment by dephosphorylating ERM proteins. Below, we discuss the different stages of this emerging pathway.

Engagement of trimeric viral Spike with dimeric host receptors via multivalent, high-avidity binding promotes receptor clustering that may generate local membrane deformation and lateral tension, well-established stimuli for PIEZO1 activation [57]. These biophysical perturbations are well-recognized triggers of mechanosensitive signaling pathways involving PIEZO1. Activation of PIEZO1 locally raises intracellular Ca^2+^ and promotes lysosome-plasma membrane fusion [16, 17]. Note that in our pathway, ACE2 clustering precedes ceramide-dependent stages rather than follows them, as suggested in [58], where ACE2 clustering depends on the formation of ceramide-enriched membrane domains. Furthermore, our finding that ASM knockout inhibits cell-cell fusion without suppressing lysosomal exocytosis (Fig. 2) argues against major contributions from other effects of lysosomal exocytosis that can promote plasma membrane remodeling by delivering lysosomal membrane to the PM and releasing other lysosomal cargo [59, 60].

Receptor clustering after ligand binding is a common mechanism of transmembrane signaling and, in some cases, has been linked to PIEZO1 activation, lysosomal exocytosis, and ASM secretion. Spike-induced ASM activation may share mechanistic features with CD95 (Fas) receptor ligation by FasL [61], and similar principles may apply to CD40, TNFR, CD28, CD2, CD44, L-selectin, and integrin activation [62]. Ligation of DC-SIGN by its natural ligand, antibodies, or measles virus also rapidly activates sphingomyelinases and induces ceramide generation at the cell surface [63].

Interestingly, a recent study identified the late-endosomal cholesterol transporter Niemann–Pick C1 (NPC1) as an additional host factor for SARS-CoV-2 entry. NPC1 interacts with the receptor-binding domain of the SARS-CoV-2 Spike protein, and Tubeimosides I–III disrupt this interaction and inhibit viral entry [64]. Notably, Tubeimoside I has also been identified as an inhibitor of the mechanosensitive cation channel PIEZO1 and was shown to antagonize Yoda1-induced PIEZO1 activation [65]. Considering our findings and the established roles of PIEZO1 in membrane mechanosensation and Ca^2+^ signaling, PIEZO1 inhibition may therefore contribute to the effects of Tubeimosides on membrane dynamics and viral entry. However, whether PIEZO1 inhibition is mechanistically linked to the disruption of Spike–NPC1 interactions or directly contributes to the antiviral activity of Tubeimosides remains to be determined.

Previous studies have reported that lysosomotropic agents, including NH4Cl [14] and niclosamide [66] inhibit syncytium formation. Given ASM’s central role in our proposed model and these agents’ ability to perturb the acidic lysosomal environment, we hypothesize that lysosomal alkalinization may weaken ASM’s membrane association, thereby altering its accessibility to lysosomal proteases. ASM dissociated from the lysosomal membrane may consequently become more susceptible to proteolytic degradation, reducing the pool of ASM available for lysosomal exocytosis and delivery to the cell surface. Dynasore, a dynamin inhibitor that also inhibits Spike-mediated syncytium formation [14], may act through a complementary mechanism by disrupting dynamin-dependent lysosomal membrane recycling [67], thereby limiting the delivery of ASM-containing lysosomes to the plasma membrane and subsequent lysosomal exocytosis. Together, these observations support a model in which both ASM stability within lysosomes and its delivery to the cell surface may contribute to efficient syncytium formation. These proposed mechanisms remain untested and will be the subject of future investigation.

When ASM is externalized, it hydrolyzes sphingomyelin on the outer leaflet of the plasma membrane, generating ceramide at the cell surface. Because PP1 and PP2A are intracellular serine/threonine phosphatases, a fundamental unresolved question is how extracellularly generated ceramide influences their activity. We propose three non-mutually exclusive mechanisms.

**In the first one**, PP1/PP2A activity is modulated by ASM-generated ceramide following rapid flip-flop of ceramide from the outer to the inner plasma membrane leaflet. Ceramides (C6, C10, C16) exhibit flip-flop half-times <1 min in model membranes at 37 °C [68]. However, ceramide generated at the cell surface does not exist as isolated monomers: ASM activity drives the formation of ceramide-enriched, gel-like domains that increase lipid packing and membrane order. Such domain-level ordering is expected to slow transbilayer movement by raising the energetic cost of desorption from the outer leaflet.

Because plasma-membrane sphingomyelin is enriched in long- and very-long-chain acyl species, ASM activity predominantly produces C16–C24 ceramides. These species may flip more slowly than short-chain ceramides due to increased acyl-chain interdigitation [69] and tighter packing. The combined effects of (i) longer acyl chains and (ii) domain-level ordering within ceramide-rich platforms may therefore substantially reduce flip-flop kinetics in living cells. Whether these constraints are sufficient to limit outer-to-inner leaflets ceramide movement—and thus restrict ceramide access to intracellular signaling pathways—remains unresolved.

**In the second mechanism**, Ca²⁺ influx through PIEZO1 activates TMEM16F [70, 71], promoting the transbilayer redistribution of ceramide and other lipids. TMEM16F also promotes the formation of SARS-CoV-2 Spike-mediated syncytia [66], supporting the possibility that TMEM16F-dependent lipid redistribution facilitates membrane fusion.

**Finally, in the third mechanism**, ASM-generated ceramide does not cross the membrane; instead, ceramide-enriched platforms initiate signaling cascades that activate intracellular nSMase. Ceramide-rich membrane domains are known to promote Src kinase activation, followed by PKC and MAPK signaling, pathways that have been implicated in nSMase activation [72]. Notably, Rottlerin, a protein kinase C-δ inhibitor, inhibited SARS-CoV-2-mediated cell-cell fusion [14]. This mechanism is consistent with our findings that nSMase activity contributes to fusion and that this contribution depends on functional ASM. Although the mechanism of transbilayer cooperation between sphingomyelinases remains unclear, our observations support a model in which ASM and nSMase function as components of a coordinated signaling axis rather than as independent enzymatic pathways.

Based on our findings and previously published studies, we propose a model for SARS-CoV-2-mediated syncytium formation in which the direct contribution of ACE2 conformational activation of Spike is complemented by Spike-induced ACE2 signaling. In this model, ACE2–Spike engagement promotes receptor clustering, generating mechanical stresses that activate the mechanosensitive channel PIEZO1. PIEZO1-mediated Ca^2+^ influx promotes lysosomal exocytosis and externalization of ASM. ASM-generated ceramide accumulates in the outer leaflet of the plasma membrane, thereby facilitating nSMase activation.

While our data indicate that Spike–ACE2 engagement promotes ASM externalization through a PIEZO1-sensitive route, and that ASM and nSMase actively support fusion, we did not directly measure ceramide or ASM activity at the cell surface. The steps connecting these observations to enhanced fusion therefore remain inferential. In addition to being an intrinsically fusogenic lipid reported to promote spontaneous fusion of protein-free bilayers of some compositions [73], ceramide is an important bioactive signaling molecule [40]. Our finding that inhibition of cell–cell fusion in ASM-knockout target cells can be rescued by activation of PP1/PP2A phosphatases and by reagents disrupting the ERM–plasma membrane linkage regulated by these phosphatases supports a role for membrane–cortex coupling in this pathway. Several of the reagents that rescued the fusion defect associated with ASM deficiency also modestly enhanced fusion in ASM-WT cells, suggesting that fusion-promoting signaling in ASM-WT cells is not fully saturated and that targeting distinct steps in the signaling pathway can further enhance fusion.

Taken together, these findings are consistent with a model in which ceramide promotes Spike-mediated syncytium formation by facilitating detachment of the actin cortex from the plasma membrane. This detachment may facilitate membrane deformation by reducing actomyosin-generated tension or attenuating its transmission to the plasma membrane.

Although weakening ERM-mediated attachment of the actin cortex to the plasma membrane promotes diverse cell–cell fusion processes, including osteoclast and viral glycoprotein-mediated fusion [50–52, 74], the specific stages of fusion facilitated by cortex–PM detachment remain to be defined.

More broadly, although the late steps in ceramide signaling and its role in SARS-CoV-2 pathophysiology are not yet fully understood, our findings place ceramide within a growing family of lipid regulators of cell–cell fusion. Several fusion processes are associated with or depend on lipid scramblase activation and non-apoptotic exposure of phosphatidylserine at the cell surface [71, 75]. Other membrane lipids, including phosphatidic acid [76], PIP2 [77], and ether lipids [78], have also been implicated in the regulation of myoblast fusion. Our findings that sphingomyelinase activation and regulation of ERM phosphorylation modulate membrane fusogenicity provide insight into how dynamic changes in membrane lipid composition and membrane–cytoskeleton interactions shape cell fusion. They may contribute to a broader understanding of fusion mechanisms and cell signaling in diverse viral and cellular systems beyond SARS-CoV-2.

## Author contributions

E.M.Z. and E.N.Z. designed and performed experiments and wrote the first draft. K.M. performed all statistical data analysis. E.M.Z, E.N.Z, K.M. and L.V.C. participated in the revision of the paper and in the discussion of results. All authors read and approved the final manuscript.

## Acknowledgements

We thank Sergei Nekhai (Howard University, Washington, DC) for providing the SMAPP1 inhibitor and Benoit Vanderpier (McGill University, Montréal, QC, Canada) for the plasmid expressing the SMPD1 gene. The research in L.V.C. laboratory was supported in part by the Intramural Research Program of the National Institutes of Health (NIH). The contributions of the NIH author(s) are considered Works of the United States Government. The findings and conclusions presented in this paper are those of the author(s) and do not necessarily reflect the views of the NIH or the U.S. Department of Health and Human Services.

## METHODS

### Cell Lines, Culture Conditions and Nomenclature

HEK293T/17 cells (ATCC, CRL-11268), parental A549 cells and A549 SMPD1-knockout cells (Abcam, ab301249) were cultured in DMEM (Gibco, 10564011) supplemented with 10% heat-inactivated FBS (Gibco, A5670801) and Normocin (InvivoGen, ant-nr-05; 1:500). For imaging-based assays, DMEM FluoroBrite (Gibco, A1896701) supplemented with 4% FBS was used to reduce background fluorescence. All cell lines tested negative for mycoplasma and were routinely monitored using the MycoFluor Mycoplasma Detection Kit (Thermo Fisher Scientific, M7006).

For clarity, we use the following designations for the cell lines employed in this study. HEK293/17 cells stably expressing SARS-CoV-2 Spike protein are referred to as HEK-Spike. Wild-type A549 cells and SMPD1-knockout A549 cells stably expressing ACE2 under the weak SV40 promoter are designated A549-ACE2-ASM-WT and A549-ACE2-ASM-KO, respectively. Both A549-ACE2 cell lines also stably express the mNeonGreen fragment mNGK1–10 and blue fluorescent protein (BFP). HEK-Spike cells additionally stably express 5×NG11 and red fluorescent protein (RFP). In selected experiments, A549-ACE2-ASM-KO cells stably expressing ASM-pHluorin were used to monitor ASM externalization, whereas A549-ACE2-ASM-WT and A549-ACE2-ASM-KO cells stably expressing LAMP1-pHluorin were used to monitor lysosomal exocytosis.

### Antibodies and Reagents

A recombinant anti-human ACE2 monoclonal antibody (Abvigen, ABSMA-2979FULL), an ACE2 Fab fragment (Abvigen, ABSMA-2979FAB), and a mouse anti-Fab antibody (Life Technologies, SA119255) were used for ACE2 binding and ligation. ACE2 (E-11) Alexa Fluor 647 (Santa Cruz Biotechnology, sc-390851) was used for ACE2 detection. SARS-CoV-2 Spike was detected using an anti-Spike antibody (CR3022; GeneTex). We used an Alexa Fluor 647-conjugated allepaca anti-rabbit IgG recombinant VHH (Chromotek, srAF647) as a secondary detection reagent. GFP fluorescence was enhanced using GFP-Booster 647 (Proteintech, gb2AF647).

All plasmids, constructs, and reagents are listed in Table 1.

**Table 1.** REAGENTS USED.

| Name of reagent | Company | cat # | Target/function | Ref. |
| --- | --- | --- | --- | --- |
| Yoda2 | Hello Bio Inc | HB9134 | PIEZO1 activator | 10.1111/bp<br>h.15996 |
| m-3M3FBS | MedChemExpress | HY-19619 | phospholipase C activator |  |
| DT-061 | MedChemExpress | HY-112929 | activator of protein<br>phosphatase 2A (PP2A) |  |
| PDP3 peptide cell<br>permeable | CEDARLANE<br>Laboratories USA Inc. | P010-610 | cell-penetrating peptide<br>activator<br>protein phosphatase-1<br>(PP1) | 10.1002/ani<br>e.20120430<br>8 |
| SMAPP1 | MedChemExpress | HY-130514 | an activator of protein<br>phosphatase-1 |  |
| GsMTx4 | GlpBio Technology Inc | GC38465 | Peptide inhibitor of<br>PIEZO1 |  |
| Blebbistatin | MedChemExpress | HY-13813 | non-muscle myosin II<br>(NMII) inhibitor |  |
| NSC305787 | MedChemExpress | HY-18931 | ezrin (Thr567)<br>phosphorylation<br>inhibitor |  |
| MMV667492 | MedChemExpress | HY-124144 | ezrin inhibitor |  |
| GSK-A1 | MedChemExpress | HY-125118 | type III phosphatidylinositol 4kinase PI4KA (PI4KIII $\alpha$ ) inhibitor | |
| nSMase2-IN-1 | MedChemExpress | HY-155111 | Neutral sphingomyelinase 2 (nSMase2) inhibitor |  |
| DPTIP | MedChemExpress | HY-131002 | neutral sphingomyelinase 2 (nSMase - 2) inhibitor |  |
| Nourseothricin Sulfate | Gold Biotechnology, Inc | N-500-1 | antibiotic |  |
| C6 NBD Ceramide (d18:1/6:0) | Cayman Chemical Co. | 62527 | lipid |  |
| C6 Ceramide (d18:1/6:0) | Avanti Research | 860506 | lipid |  |
| GFP-Booster 647 | Proteintech Group | gba647n | GFP-binding Ab |  |
| Spike Glycoprotein (Stabilized) from SARS-Related Coronavirus 2, Wuhan-Hu-1 | BEI RESOURCES.ORG | NR-53524 | Spike Glycoprotein (Stabilized) from SARS-related Coronavirus 2, Wuhan-Hu-1 |  |
| mVenus N1 plasmid:SMPD1 | Gift from Benoît Vanderperre |  |  | 10.1002/m<br>ds.27642 |
| Lenti SMPD1-SEpHluorin |  |  | SMPD1-Venus was replaced by SMPD1-SEpHluorin | This work |
| Lenti Lamp1-SEpHluorin |  |  | Lamp1-GFP was replaced by LAMP1-SEpHluorin | This work |
| Lentivirus pLEX307-ACE2 Blast | Addgene | #158449 | ACE2 expressing Lentivirus | This work |
| Lenti SV40-ACE2 Ef1a-Blast |  |  | SV40-ACE33 Expressing Lentivirus | This work |
| pCMV14-3X-Flag-SARS-CoV-2 S | Addgene | #145780 | SARS-CoV-2 Spike |  |
| Lenti SMPD1-EF1aHygro |  |  |  | This work |
| <u>pRSV-Rev</u> | Addgene | 12253 |  | 10.1038/s41467-020-15562-9 |
| pMD2.G | Addgene | 12259 |  |  |
| LEGO G/Bsd | Addgene | 27354 |  |  |
| psPAX2 | Addgene | 12260 |  |  |
| Red FP | Addgene | 74252 |  |  |
| mTagBFP2 | Addgene | 113725 |  |  |

### Production of lentiviral particles

Lentiviral particles were generated in HEK293T/17 cells by co-transfecting the packaging plasmids psPAX2, pMD2.G, and pRSV-Rev with the transfer plasmid encoding the gene of interest. The plasmids were combined at a 1:1:0.5:2 ratio, diluted in Opti-MEM (Gibco, 51985034), and mixed with Hieff Trans Booster DNA/RNA Transfection Reagent (Yeasen, 40801) according to the manufacturer’s instructions. After 6–8 hours, the medium was replaced with DMEM supplemented with 10% FBS. Cells were incubated at 37°C for 48 hours. Supernatants containing lentiviruses were collected, cleared through 0.45 µm filters, and used for two weeks.

### Generation of ACE2-Positive A549-ACE2-ASM-WT and A549-ACE2-ASM-KO Clones

A549-WT and A549 ASM-KO cells were transduced with lentiviral vectors encoding human ACE2 under the control of a weak SV40 promoter and selected under antibiotic pressure for 1 week. The details of the transduction procedure are published in https://www.addgene.org/protocols/lentivirus-production. Following selection, the cells were passed twice without antibiotics. Individual clones were screened to identify matched ACE2 surface expression levels in the WT and KO backgrounds. Selected clones were expanded and subsequently transduced with lentiviruses encoding mNG3K1-10 and a blue fluorescent protein.

### Selection of HEK293T/17 Spike-Expressing Clones

HEK293T/17 cells were transduced with lentivirus encoding SARS-CoV-2 Spike (Wuhan strain, wild type) and subsequently with lentiviruses encoding 5xNG11 and a cytoplasmic red fluorescent protein. Following antibiotic selection, the clones were stained with an anti-Spike antibody at 1:500. An Alexa Fluor 647-conjugated alpaca anti-rabbit IgG recombinant VHH was used as a secondary detection reagent at 1:1000.

Clones exhibiting robust Spike surface expression were expanded under antibiotic pressure and subsequently passed twice without antibiotics before experimentation.

### Fusion Assay Using HEK293-Spike and A549-ACE2 Cells

A549 cells expressing ACE2, mNG3K1-10, and the cytoplasmic blue fluorescent protein x2mtagBlue2 (A549-ACE2-ASM-WT) or (A549-ACE2-ASM-KO) were seeded onto 18-well Ibidi slides (81816) and cultured to 85–90% confluence. HEK293-Spike cells expressing 5xNG11 and the cytoplasmic red fluorescent protein Ruby3 were detached using Cellstripper (Corning, 25-056-01), resuspended in complete medium, and added to the A549 monolayers in the presence or absence of the indicated reagents. The red and blue fluorescent proteins enabled clear discrimination between donor and target cells, assessment of their uniform distribution, and visual monitoring of cell after pharmacological treatments. Cells were incubated for 2 hours at room temperature to facilitate heterotypic cell contacts, followed by incubation for 36 hours at 37°C. The reagent concentrations (Figure legends) were determined experimentally by prior titration to maximize treatment effect without compromising cell viability, which was identified by DAPI staining (1 µg/mL; Thermo Fisher Scientific, D21490) or the LIVE/DEAD Cell Imaging Kit (R37601)

After incubation, live cells were imaged using a Zeiss LSM 800 confocal microscope. The entire well in an 18-well plate was imaged. Syncytia were identified through mNeonGreen complementation. Image quantification was performed using Fiji ImageJ (v1-54p). Individual syncytia were detected as follows: background signal was removed using the rolling ball algorithm with a 20-pixel radius, images were smoothed with a Gaussian kernel of 2-pixel radius, and thresholding was applied using the Otsu method. The area of each syncytium was measured using the Analyze Particles command. Efficiency of fusion was quantified as the average area of syncytia for each experiment. Each experiment was conducted 6 to 7 times.

### Measurement of ASM Externalization

A549-ACE2-ASM-KO cells were stably transduced with lentiviral vectors encoding ASM-pHluorin and selected using antibiotics, as described above. The cells were subsequently transduced with a lentiviral vector encoding a GPI-anchored red fluorescent protein to facilitate visualization of the plasma membrane.

A549-ACE2-ASM-KO cells stably expressing ASM-pHluorin were treated for 4 hours at room temperature (to minimize internalization) with stabilized trimeric SARS-CoV-2 Spike (10 µg/mL), Yoda2 (5 µM), or a combination of Spike and GsMTx4 (12 µM). Cells were fixed for 15 minutes at 37°C by adding an equal volume of 8% paraformaldehyde, yielding a final concentration of 4%. The cells without permeabilization were then washed with PBS, blocked with 5% goat serum, incubated with GFP-Booster 647 (1:200) for 1 hr. at RT, washed with PBS, and analyzed.

In parallel experiments, A549-ACE2-ASM-KO stably expressing ASM-pHluorin were treated for 4 hours at room temperature with an ACE2 antibody (20 µg/mL), an ACE2 Fab fragment (7 µg/mL), or the Fab fragment combined with an anti-Fab antibody (30 µg/mL). Cells were then fixed without permeabilization and stained with GFP-Booster 647. As a control, the cells that were not transduced with ASM-pHluorin were treated with GFP-Booster 647 under the same experimental conditions as above.

We imaged 15 fields, each containing 6-7 cells. Background was removed using a rolling ball algorithm with a 20-pixel radius, and images were smoothed using a Gaussian kernel with a radius of 2 pixels. Fluorescent puncta corresponding to externalized pHluorin were identified by thresholding with the Otsu method. We calculated the total intensity of all fluorescent puncta in each field and averaged these values across all fields. Each experiment was performed at least four times.

### Measurement of LAMP1-pHluorin Externalization

A549-ACE2-ASM-WT cells and A549-ACE2-ASM-KO cells (both lines stably expressing LAMP1-pHluorin) were untreated or incubated with Yoda2 (5 µM) for 4 hours at room temperature, then fixed without permeabilization and stained with GFP-Booster 647, imaged, and analyzed as above.

### Statistical Analysis

Data were analyzed in R (v4.6.1) using linear models. For experiments with a single cell line, we used the formula log(Y) ∼ Treatment; for experiments with multiple cell lines, we used log(Y) ∼ CellLine * Treatment. The response was log-transformed to address heteroscedasticity across factor levels. Y axis on all plots of cell fusion quantification is log transformed to match our statistical model. Pairwise comparisons (two-sided) were performed using the emmeans package (v2.0.4), with p-values adjusted for multiple comparisons using the Šidák method. Significance is denoted by asterisks (*p < 0.05, **p < 0.01, ***p < 0.001; ns, not significant) in the figures.

